# Novel method to assess group dynamics in rats reveals deficits in behavioral contagion in KM rats

**DOI:** 10.1101/2024.08.30.610455

**Authors:** K. Smirnov, I. Starkov, O. Sysoeva, I. Midzyanovskaya

## Abstract

Behavioral copying is a key process in group actions, but it is challenging for individuals with Autism Spectrum Disorder (ASD). We investigated behavioral contagion, or instinctual replication of behaviors, in Krushinky-Molodkina (KM) rats (n=16), a new rodent model for ASD, compared to control Wistar rats (n=15). A randomly chosen healthy Wistar male (“demonstrator rat”) was introduced to the homecage of experimental rats (“observers”) 10-14 days before the experiments to become a member of the group. For the implementation of the behavioral contagion experiment, we used the IntelliCage system, where rats can live in a group of 5-6 rats and their water visits can be fully controlled. During the experiment, the demonstrator was taken out of IntelliCage for 24 hours of water deprivation and then placed back. As a result, a drinking behavior of the water-deprived demonstrator rat prompted activated behaviors in the whole group. Unlike the Wistar controls, KM observers showed fewer visits to the drinking bottles, particularly lacking inspection visits. The control group, in contrast, exhibited a dynamic, cascade-like visiting of the water corners. The proportion of activated observers in KM rats was significantly lower, as compared to Wistar ones, and they did not mimic other observer rats. KM rats, therefore, displayed an attenuated pattern of behavioral contagion, highlighting social deficits in this ASD model. This study suggests that measuring group dynamics of behavioral contagion in an automated, non-invasive setup offers valuable insights into social behavior in rodents, particularly for studying social deficits in ASD models.

**Highlights:**

- Thirsty demonstrators triggered an avalanche of observers’ visits to the water corners
- The contaged behavior was attenuated in observer KM rats
- Behavioral contagion test provides a new tool for objective, automated phenotyping in rodent models of social deficits

## 1. Introduction

Behavioral copying is the simplest mode of social learning where a mood, attitude, or behavior spreads quickly from one individual to another (Polansky, Lippitt and Redl, 1950). This phenomenon enhances coordination within a group and fosters social cohesion, also reducing mutual aggression (Laméris *et al*., 2020). However, this mode of behavior is deficient in individuals with Autism Spectrum Disorder (ASD) (Stewart, McIntosh and Williams, 2013). The decreased ability to replicate the behavior of other group members leads to failures in social or academic training (Pittet *et al*., 2022) necessitating specialized tutorial support for ASD pupils. The development of animal models for ASD is crucial for advancing treatment strategies, similar to those seen for other neurodevelopmental and neurodegenerative diseases (Li *et al*., 2024; Pan *et al*., 2024). Animal models allow experimental approaches to study social deficits and, in translational perspectives, lead to more effective support for ASD patients (Chadman, Guariglia and Yoo, 2012; Ruhela, Prakash and Medhi, 2015; Sierra-Arregui *et al*., 2020; Silverman *et al*., 2022).

Recently, we proposed that the Krushinsky-Molodkina (KM) rat strain, originally bred as a model for convulsive epilepsy (rev. in Poletaeva *et al*., 2017), may also serve as an animal model for ASD (Rebik *et al*., 2022, 2023). In various experimental contexts, KM rats showed consistent deficits in social contact motivation. These social deficits were observed both in seizure-naive KM rats and in KM rats with moderate seizure experience (Rebik *et al*., 2023). Besides the ASD-like behavioral phenotype, KM rats also showed imbalanced binding to D1-like and D2-like dopamine receptors in the insular cortex (Birioukova, Van Luijtelaar and Midzyanovskaya, 2024), a region also dysfunctional in ASD patients (Mittleman and Blaha, 2015; Sato *et al*., 2023; Blum *et al*., 2024). Thus, the KM rat model shares not only an ASD-like phenotype, but also a part of the pathophysiology known for ASD.

We hypothesized that engagement in collective actions would be difficult for KM rats compared to healthy rats. To reduce environmental factors affecting social behavior, such as anthropogenic stress and novelty-induced environmental dishabituation, we adapted the behavioral contagion approach previously reported for dyadic interaction in rodents (Ivanov, Semenov and Krupina, 2014). In those studies, a demonstrator rat, isolated and water-deprived for 24 hours, started drinking upon reintroduction, and this behavior was replicated by a non-deprived observer rat (Ivanov, Semenov and Krupina, 2014; Ivanov and Krupina, 2017). This contaged behavior was absent if the demonstrator rats were not motivated (i.e., not water-deprived) (Ivanov, Semenov and Krupina, 2014).

In this study, we report a modified paradigm of behavioral contagion to be automatically tested in groups of normotypic and autistic phenotype rats. The experiment was conducted in an IntelliCage setup, housing 4-5 animals simultaneously (Lipp *et al*., 2024). The drinking bottles were located in corner compartments, where access to water was limited up to by automated doors. This IntelliCage setup allowed for the identification of each individual animal inside the drinking corners. The demonstrator rat, which had been a group member for at least 10-14 days before the cohort entered the IntelliCage, was isolated and water-deprived for 24 hours (Fig. 1A). Soon after reintroduction, the thirsty demonstrator rats started drinking, while the intact observer rats behaved freely and copied (or did not copy) the water corner visits. We hypothesized that the contaged response would be attenuated in KM rats due to their deficient social motivation (Rebik *et al*., 2022, 2023).

**Figure 1.**
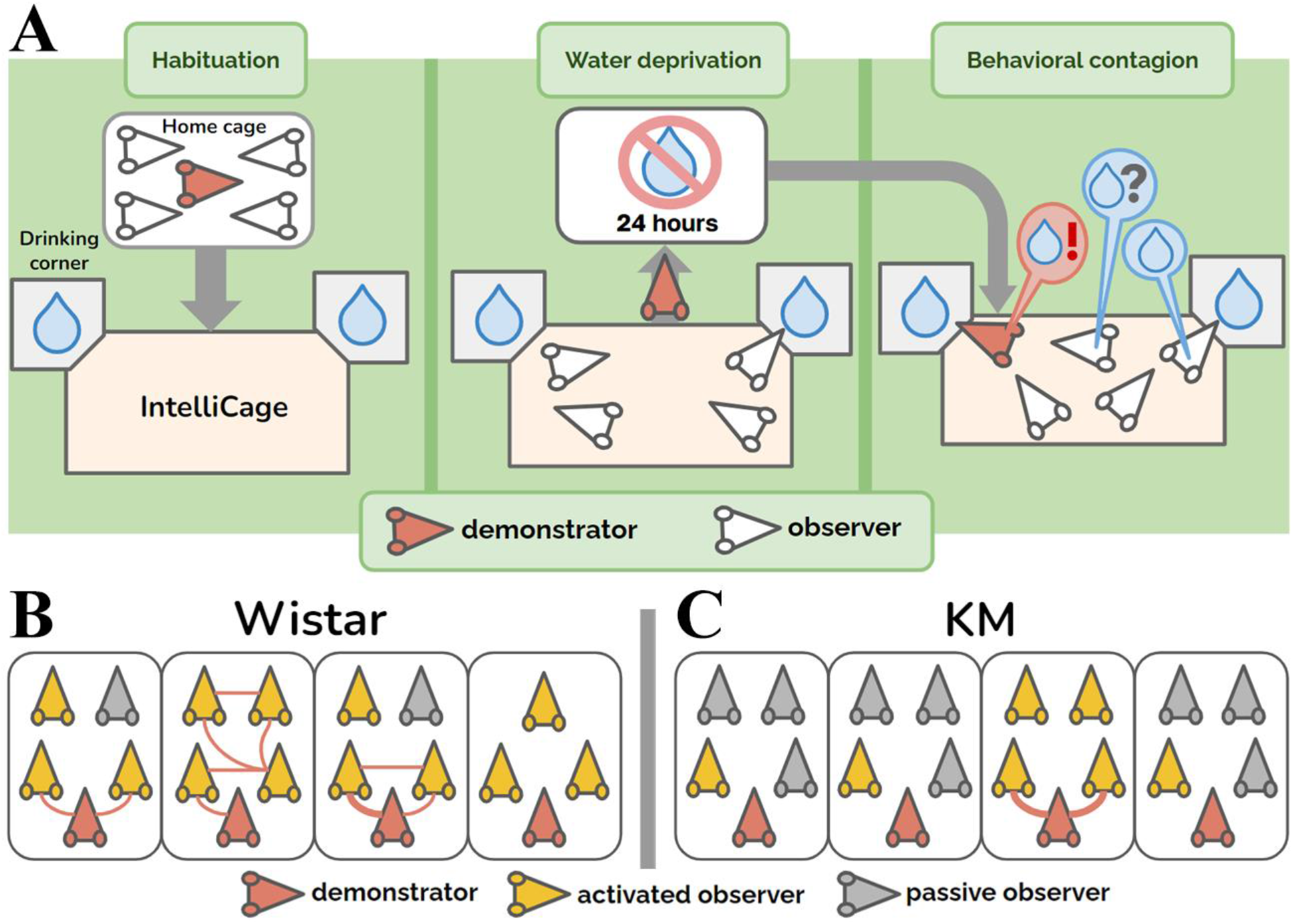
Group aspects of behavioral contagion. A: Experimental design. B, C: Group graphs in cohorts of control Wistar (panel B) and KM rats (panel C). The group graphs are drawn for the 30 minutes of observation. The red rat heads mark the demonstrator (i.e., water-deprived) rats in each group. The yellow rat heads denote the activated observer rats, i.e., the non-deprived animals that made at least one visit to a water corner. The gray rat heads correspond to the passive observers, which did not make any visits to the water corners. The red edges between the nodes mark fast (<4s) copied visits to a water corner, demonstrated by any groupmate, with the line thickness proportional to the number of copied visits (ranging from 0 to 3 in our study).

## 2. Methods

### 2.1 Animals

The experiments enrolled intact adult male rats, aged between 6 and 8 months, and weighing 350-450g. The cohorts were 15 outbred Wistar rats and 16 inbred KM rats. The animals arrived at the animal chapter of IHNA at the age of 6-8 weeks. The rats were housed 4-5 in a cage, with free access to water and standard pellet food, under 12h light/dark regime (lights on at 8:00 A.M) A month prior to the main experiment, all the rats were implanted with individual identification veterinary chips below the skin and above the neck muscles under mild sedation induced by intramuscular injection of dexmedetomidine (0.2 mg/kg). A new, randomly chosen and previously unfamiliar Wistar male rat of the same age and weight was added to each group of 3-4 rats, 10-14 days prior to the main experiment to serve as the demonstrators later on. The given time period of being housed in a group with rats of a different strain is sufficient for the albino rat to display pro-social behavior towards rats of the other strain (Ben-Ami Bartal *et al*., 2014). After the given period of common housing, the cagemates were considered as a group in our further experiments.

### 2.2 IntelliCage Set-Up

We used the IntelliCage set-up with two drinking corners available to the rats. The pre-implanted veterinary chips allowed collecting individual information on the entries, occupation time and the number of lickings made inside the water corners.

The experiment in the IntelliCage setup consisted of three blocks (Fig.1A). All the procedures, i.e. the start of all blocks, water deprivation protocols, and re-introduction of the individual demonstrator rats, were set up at 12.01 PM. For the first three days, the animals adapted to the experimental set-up. The doors to the drinking bottles were opened every time a rat was detected inside a corner. The drinking time was unlimited. Then the second adaptation block followed, during which the rats were trained to open the doors by poking their noses into a special area inside the corners. The door opening time was limited to 15 seconds, after which the rat had to make a new entry in order to access water again. The third block was the water-deprivation procedure. The demonstrator rat was withdrawn from the group, put away in the home cage without a drinking bottle available, and the cage was placed in another room. All the rats left in the IntelliCage (observers) had the same conditions of water access, as earlier. For the demonstrator rat, water deprivation lasted for 24 hours. After this, the demonstrator rat was returned to the group mates. The activity registration started by the time of the demonstrator re-introduction, and lasted for at least 4 hours. The deprived rat demonstrated the most pronounced drinking behavior within the first hour (Fig.2) after re-introduction. The time interval of 30 minutes was chosen for analysis of behavioral contagion.

### 2.3. Experimental registration

Each water corner had the sensors for the rat veterinary chips, and two drinking bottles equipped with the sensors of the lick contact times. The following parameters were registered for each water corner’s approach: the rat ID, the times of entry and leaving, and the number of licks made during the visits.

### 2.4. Graph construction

The behavior of grouped rats was presented as a graph, built up for the 30 minutes of behavioral contagion, for each individual group (Fig 1B, C). Several definitions were accepted to construct the graph representations:

- For each graph, the nodes were individual rats, and the edges’ thickness represents the association index (i.e. the number of copied visits).
- A water corner was considered as the demonstrated one, immediately after a visit of any rat in the group.
- The visit was regarded as a copied one (red lines as the graph edges, Fig. 1B), if a rat entered the demonstrated water corner with a latency <4s. The threshold was modified from a previous work on a contagious water consumption (Ivanov *et al*, 2014), where it was equal to 2s. The latency of copied visits was enlarged here due to a complex geometry of the IntelliCage’s water corners, with its tunnel-like entrance chambers, preceded by a step.
- An observer rat was considered as an activated one, if it visited any water corner (yellow rat head, Fig 1B, C) during the behavioral contagion window. Otherwise, the rat was depicted as a passive observer (gray rat head, Fig. 1B, C).

### 2.5. Statistics

In this study, we used a two-factor repeated measures ANOVA, for the following parameters: the total number of water corner visits, the number of drinking visits, the number of inspection (i.e., without drinking) visits, and the total time of licking. All the data were time-normalized. Since the behavioral contagion was assessed within a 30-minute interval, the baseline data, collected for three hours, was divided into 6. The strain was taken as a “between” factor; the time conditions (baseline and contagion) were considered as a “within” factor. In cases of significant interaction of the factors, post-hoc Bonferroni tests were used to assess the differences. We also considered correlations (all were done using Spearman’s method) between different metrics of the rat groups. Namely, we studied whether the pre-contagious activity would affect the individual response to the demonstrator behavior. The individual pre-contagious activity (licking numbers, drinking entries, inspection entries) had been collected for 3 hours, 8.50-11.50 a.m., and correlated with the same parameters, as measured for the 30 minutes of contagious behavior.

The assessment of collective activity was carried out on the basis of individual group graphs (Fig. 1B, C). The proportion of active (i.e., who made at least 1 visit during 30 minutes of collective contagion) observers was compared between the Wistar and KM cohorts using Chi-square test (Tables 2*2). The same was done for the proportion of copying from observer to observer within the groups: the actual number of red edges between yellow “rat head” signs of activated observers, referring to the maximum possible number of all connections between all observers (which is (N^2^-N)/2, where N is the group size). The group parameters (the number of actual edges and the number of all possible edges) are summarized for each strain and compared using Chi-square test for the resulting 2*2 tables.

## 3. Results

### 3.1 The behavioral contagion phenomenon

Behavioral contagion was expressed as an avalanche of simple acts, including water corner visits and water consumption, registered in the rats. The observer rats, being intact and non-deprived, responded (or did not respond) to the demonstrator’s agitated drinking behavior (Fig. 2A, B). All the water-deprived (demonstrator) rats began their serial drinking attempts as soon as they were reintroduced to the IntelliCage. The demonstrators continued shuttling between the available water bottles for a prolonged period, as seen from the experimental timelines (upper two lines on each chart, shaded with gray on Fig. 2A, B).The time interval of 30 minutes was chosen for analysis of behavioral contagion, to catch the most probable period of groups’ responses (Fig2A).

**Figure 2.**
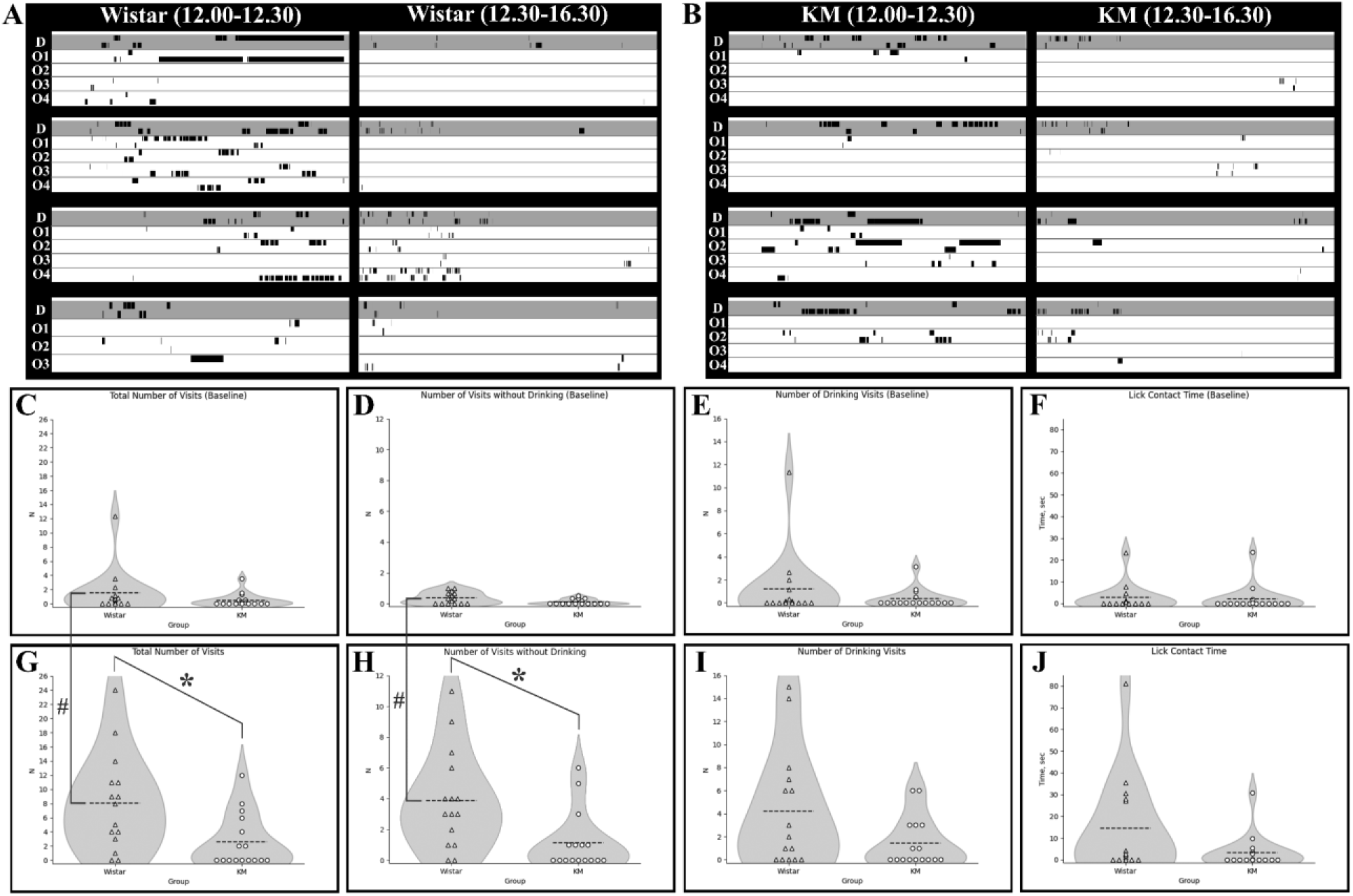
Behavioral contagion in Wistar and KM rats. A, B: The water corner visits were recorded in Wistar (WS, panel A) and KM (KM, panel B) rats. The activities are shown for the first 30 minutes (left panels) and the following 4 hours (right panels), starting from the re-introduction of demonstrators (the first two lines in each chart) to their group mates. Each pair of striped lines corresponds to an individual rat (underlined); the timelines represent visits to drinking corners 1 (the upper rows of stripes) and 2 (the lower rows of stripes). The thickness of the black stripes is directly proportional to the occupation time for a water corner. C-J: Inter-(* - p<0.01) and intragroup (# - p<0.01) differences. C: Total number of visits during baseline. D: Number of inspection visits during baseline. E. Number of drinking visits during baseline. F. Drinking time during baseline. G: Total number of visits during behavioral contagion phase. H. Number of inspection visits during behavioral contagion phase. I: Number of drinking visits during behavioral contagion phase. J: Drinking time during behavioral contagion phase.

### 3.2 Baseline activity of Wistar and KM rats preceding behavioral contagion

To assess the baseline behavior of the animals, we selected a time period from 8:50 to 11:50. This period started 50 minutes after the lights were turned on, allowing the rats to adapt to the change in lighting, and ended 10 minutes before the demonstrator’s return, as the experimenter might enter the room with the setup a few minutes before 12:00. The three-hour time period was chosen because rats rarely make spontaneous visits to the drinking corners during the light phase, and selecting shorter time periods would increase random fluctuations of the parameters. We found no differences between the Wistar rats and KM rats in terms of the number of animals? activated (making visits) during the baseline period (8:50 to 11:50 a.m, 10 out of 15 Wistar rats, 7 out of 16 KM rats; Chi-square = 1.643, p = 0.199), nor in the total number of visits {F(1,30)=8.97, p=0.19}, the number of drinking visits {F(1,30)=5.26, p=0.29}, and the duration of drinking {F(1,30)=2.99, p=0.78}.

### 3.3 Strain Differences in Behavioral Contagion

The control rats demonstrated dynamic engagement of group members in behavioral contagion (Fig.1B), which was poorly observed in KM rats (Fig. 1C). Specifically, the proportion of activated observers (i.e., group members that made at least one visit to a water corner during the contagion period, yellow rat heads on Fig. 1B,C) was significantly lower in the KM cohort (7/16 in KM rats vs. 13/15 in Wistar rats; Chi-square = 6.229, p=0.012). Additionally, the proportion of followers (i.e., group mates which replicated at least one visit, red links in Fig. 1B,C) was also significantly lower in the KM cohort compared to the Wistar cohort. Specifically, 2 out of 16 KM observer rats made copied visits to the demonstrated water corners, compared to 8 out of 15 Wistar observer rats (Chi-square = 5.907, p=0.016); or, in other terms, KM rats actualized 2 out of 38 possible edges within their 4 groups, and Wistar rats actualized 10 out of 36 possible edges within their 4 groups (Chi-square=7.393, p=0.007 for the strain difference). The groups of Wistar and KM observers differed in their response to the re-introduced demonstrator cagemate. Both the “strain” and “conditions” were significant factors for all types of visiting activity. The Wistar cohort displayed higher values of the above-mentioned parameters, and also the test condition (i.e., behavioral contagion, with baseline and contagion taken as repeated measures) led to significant facilitation of the visits (Fig.2 C-J). Namely, the effects of “strain” and “conditions” were the following: the total number of visits ({F(1,29)=9.24, p=0.005} for “strain”; {F(1,29)=15.28, p=0.0005} for “condition”; the interaction of “strain”*”condition”{F(1,29)=3.87, p=0.06}, respectively), the number of drinking visits ({F(1,29)=5.26, p=0.03} for “strain”, {F(1,29)=6.68, p=0.01} for “condition”, insignificant interaction), as well as for the inspection visits ({F(1,29)=9.66, p=0.004} for “strain”, {F(1,29)=23.83, p=0.0003} for “condition”; the interaction of “strain” * “condition” significant: F(1,29)=7.26, p=0.01). For the total number of visits and for the number of inspection visits, the post-hoc Bonferroni tests were applied, due to the factors interaction. It appeared that the Wistar observer displayed a pronounced behavioral activation, unlike that seen in KM rats. Namely, the total number of visits was significantly higher in the WS groups compared to the KM groups (Fig. 2C; p=0.005, Bonferroni test). Closer analysis revealed that this difference was primarily due to facilitated inspections (Fig. 2D; p=0.0008, Bonferroni test), rather than drinking visits (Fig, 2I, p=0.10).

Behavioral contagion was not correlated with the baseline water consumption in observers of both rat strains (according to the Spearman rank order correlation, all p’s>0.10), indicating the independence of the induced behavior from putative drinking motivation. Additionally, no correlations were found between other baseline measures and behavioral parameters measured during behavioral contagion (all p’s>0.10).

To summarize, KM rats displayed a remarkably attenuated pattern of behavioral contagion, with a low number of mimicked behavioral acts, expressed by a few activated observer rats.

Therefore, the rats with ASD-like phenotype were characterized by a poor group engagement into the demonstrated behavior, unlike that seen in control rats. Normal Wistar rats were activated by the demonstrator’s agitated movements, replicating the entries to the water corners after the demonstrators, as well as after the other group members. The copied visits occurred immediately, and also with some latencies within the registered contagion period.

## 4. Discussion

### 4.1 Components of behavioral contagion

Engagement in group behaviors is a basic survival mechanism in social animals, essential for activities like social foraging and mobbing. In foraging, it is more efficient to join group success rather than making random independent choices. In this study, we observed that the simplest mode of group action— mimicking of a behavioral pattern —is well expressed in laboratory rodents, outbred normotypic Wistar (WS) rats.

Behavioral contagion is a form of allelomimetic behavior that plays an important role in the cohesion and synchronization of behavior within a social group of animals (Ginelli *et al*., 2015). According to L. Wheeler (Wheeler, 1966), behavioral contagion should be distinguished from conformity, imitation, social pressures, and social facilitation. While “social conformity” and “social pressure” are challenging to assess in animal experiments, “imitation” (behavioral mimicking) and “social facilitation” are valid concepts. Observer rats responded to demonstrators’ behaviors both by drinking and by inspections of water corners.

Further experiments are required to discriminate between imitation, goal emulation, and social facilitation as reasons for contagious behavior in normotypic rats. Notably, there was no correlation between the amount of water consumed during the baseline period and the number of water corner visits, indicating that intrinsic motivation to drink was not necessary for observers to copy the demonstrators’ behavior. This allows us to distinguish the observed phenomenon from social facilitation, which leads to the activation of motivated behavior while observing its execution in another subject (Redd and de Castro, 1992).

In the dyadic paradigm of behavioral contagion, thirsty demonstrators prompted observer rats to attend the water bottles more frequently, as compared to unmotivated observers, with this contagion lasting several minutes (Ivanov, Semenov and Krupina, 2014). Here, we set up a group test that required extending the observation period because of a larger number of participating animals and interactions between the observers. Motivated behavior consists of behavioral sequences organized within nested states of action (Sanabria *et al*., 2019). The modified behavioral contagion test allowed us to distinguish between visits made for water consumption and those made only for inspecting the bottles.

Social foraging implies that each animal watches the foraging success of a group member to share the found resource. We observed that control rats were more likely to inspect visited water bottles rather than drink from them (Fig. 2G, H, I). Active attention to the behavior of a group member, expressed here as inspections of recently visited water corners, is essential for social learning in gregarious animals. Attention to conspecifics’ behavior is a prerequisite for contagious yawning, another model of neutral behavior that spreads in animal groups (Gallup, 2021).

### 4.2 Attenuated behavioral contagion in KM rats

As hypothesized, KM rats showed poor reactions to demonstrators’ behavior, with low individual responses and group engagement. This extends our previous findings on deficient social motivation in KM rats (Rebik *et al*., 2022, 2023). The present setup allowed observer rats to behave freely, minimizing stress and providing an objective way to test social deficits in rodent models of ASD.

Attention may play a crucial role in the contagious inspection visits observed (Gallup, 2021, 2022). Rats with an ASD-like phenotype may pay less attention to the demonstrated actions, resulting in minimal motor replication. Special training approaches improve motor imitation in children with ASD, aiding social coping (Paparella and Freeman, 2021). Motor imitation in early childhood enhances language abilities, while reduced observational learning in adults with ASD-like traits is linked to reduced goal emulation (Wu *et al*., 2024). Imitation attempts in adults with ASD are part of social camouflage (Alaghband-rad, Hajikarim-Hamedani and Motamed, 2023). In rodents, social camouflaging is unlikely, but insufficient imitation abilities are detectable. In adult KM rats, deficiencies are seen at the level of behavioral imitation. Special experiments are needed to infer the ability of goal emulation in KM rats. Experimental manipulations that facilitate social learning in rodent models of ASD may serve as the preclinical stage for future research in ASD therapy.

The level of individual behavioral copying in animal models for ASD can serve as a translational point for basic research in ASD neurobiology.

### 4.3. Group dynamics

Group aspects of the behavioral contagion test offer new ways to study sociability in laboratory rodents, which are highly gregarious species (Latané, 1969). Rats sharing the same housing enclosures form non-random huddling associations, demonstrating established social relations (Proops *et al*., 2021). Despite recognizing the lack of knowledge on rat social interactions in the mid-20th century (Davis, 1953), methodological limitations persist, as most experiments remove animals from their usual conditions for short-term testing. Recent experimental systems allowing long-term and automated behavior registration have highlighted the complexity and importance of group interactions in laboratory rats (Nagy *et al*., 2023).

Studies on interaction networks in schooling fish reveal small groups of strongly connected neighbors who are both most socially influential and most susceptible to social influence (Rosenthal *et al*., 2015). In bird flocks, the phenomenon of murmuration demonstrates how the behavioral state of one bird influences, and is influenced by, all other birds in the event (Cavagna *et al*., 2010). While individual actions within a group are essentially stochastic, assessing intrinsic group dynamics aids in understanding behavioral contagion (Rosenthal *et al*., 2015).

The rat groups in this study were smaller than flocks of fish or birds, but dynamic behavioral effects were still observed. In normotypic Wistar rats, activated members served as a demonstrator for others (Fig. 1B), facilitating social action. Non-deprived observer rats replicated water corner visits, even with no need for additional drinking.

In contrast, the “autistic” KM cohort showed poor socially mediated engagement (Fig. 1C). KM group members did not mimic each other and showed minimal response to demonstrators. It is unclear whether the demonstrators failed to gain a high social rank in the KM cohorts during the preliminary common housing period. While one might hypothesize that low-ranked members are not followed, this should not be a strict rule for foraging-related behaviors like water consumption. Success sharing in foraging should be effective regardless of group rank (otherwise, it decreases the chance to find food resources). Additionally, KM observers did not copy each other within their subgroups (Fig. 1C), suggesting that social rank is unlikely to strongly affect contagion effects, although future experiments are needed to clarify this. It is important to note that low locomotion is a behavioral trait of KM rats (Rebik *et al*., 2022, 2023), which might contribute to their poor ability to rapidly replicate behaviors (depicted here as red edges in Fig. 1B,C). However, over the 30-minute period analyzed, the group dynamics in KM observers differ from those in the control group (indicated by gray and yellow rat heads in Fig. 1B,C). Even the slowest animals had the opportunity to visit at least one water corner during this time. KM rats, as a rule, did not do this, resulting in a low number of activated observers (evidenced by the low proportion of yellow versus gray rat heads in Fig. 1C). This suggests that slow locomotion is unlikely to be the primary reason for the poor behavioral contagion response observed in KM rats.

### 4.3. Lack of non-aversive tests for behavioral contagion

Most experiments studying behavioral contagion use emotionally negative procedures, such as painful stimuli or stressful environments, applied to demonstrators (Carnevali *et al*., 2017; Dimitroff *et al*., 2017; Qu *et al*., 2023; Auer *et al*., 2024). Stressful experiences are transmitted among conspecifics through ultrasonic vocalizations (Brudzynski, 2013; Wöhr *et al*., 2015; Bruder, Stewart and McGrath, 2017), olfactory stimuli (Kiyokawa *et al*., 2009; Takahashi *et al*., 2013; Raynaud *et al*., 2015), and other social interactions (Carnevali *et al*., 2017). Contagious yawning (Platek, Mohamed and Gallup, 2005; Massen and Gallup, 2017; Gallup, 2022) is a rare example of spreading an emotionally neutral spontaneous behavior in animals. Thus, there is a evident lack of emotionally neutral experimental setups for studying social facilitation of learning and habituation.

A remarkable feature of our approach is its non-aversiveness. This experimental paradigm could be applied to study phenomena such as social learning and intragroup hierarchy. Social learning and behavioral imitations are particularly deficient in patients with ASD. Negative emotional paradigms are often ethically unacceptable for research in human participants with ASD. So, emotional neutrality makes this and similar paradigms well translatable for preclinical tests in rodent models of ASD.

### 4.4. Limitations and further directions

As a limitation of our study, one might point to a lack of any behavioral registrations beyond the water corners. Unfortunately, the standard IntelliCage settings do not allow the registration of behaviors, such as locomotion, vocalization, grooming, aggression, or play. Social interactions involve emotional exposure (Herrando and Constantinides, 2021). Emotional contagion is crucial in conspecific interaction, facilitating social learning, empathy, and group actions (Keysers *et al*., 2022; Pérez-Manrique and Gomila, 2022). Since deficient activation and behavioral copying were observed in the KM rat cohort, new experiments are needed to study the emotional states of the observers and demonstrators.

Other psychosocial disorders, like self-harm or suicidal behavior, might have a contagious component (Seong *et al*., 2021). Therefore, a new quantitative approach to studying behavioral contagion in animals will be beneficial for preclinical and basic research beyond ASD neurobiology.

The usage of only male rats, and the employment of only normotypic demonstrator rats are the other limitations of the study. The female KM rats are not available from the breeder (Biological Faculty of Moscow State University), so the comparison of female individuals of Wistar and KM strains was not planned. Further experiments are needed to clarify details of the drinking-related behavioral contagion within female rats groups. We used only normotypic (Wistar) demonstrator rats, to have the same conditions for the comparison of observers. It is not clear what would be the impact of “autistic” demonstrators on behavior of their group mates. It is quite possible that the individuals lacking social motivation would be outcasted by other group members. However, new experiments are warranted to study this point in sufficient detail.

## Conclusion

This study introduces an emotionally neutral paradigm for investigating behavioral contagion. Using the IntelliCage setup, we observed that normotypic non-water deprived Wistar rats mimicked the water-deprived conspecifics, visiting the drinking corners. Moreover, they started to follow each other unmotivated visits, peaking within 30 minutes of water-deprived demonstrator introduction. At the same time, KM rats showed significantly reduced behavioral contagion, as compared to normotypic Wistar rats. This confirms their deficient social motivation and behavioral imitation.

Our novel approach, utilizing non-aversive experimental paradigm of behavioral contagion, offers a robust framework for exploring social behaviors in rodent models, aiding in the understanding and development of therapeutic strategies for ASD and other psychosocial disorders.

## Competing interests

The authors declare that they have no competing interests.

## Author contributions

K.S. conceived and designed the study, performed experiments, collected data, analyzed data, interpreted results, and wrote the manuscript. I.S. contributed to data analysis. O.S. contributed to result interpretation and manuscript writing. I.M. suggested using KM rats as a model of ASD, contributed to data analysis, result interpretation, and manuscript writing. All authors read and approved the final version of the manuscript.

## Funding

The work was partly supported by RSF, grant No. 23-25-00484

## Institutional Review Board Statement

The animal study protocol was approved by the Institutional Ethics Committee of IHNA and NPh RAS, (protocol No. 01.25.02.20021 from 25 February 2021)

## Acknowledgments

The authors are deeply grateful to Dr. N.A.Krupina for inspiring the study, and for invaluable scientific debates. P.L. Aleksandrov kindly helped to resolve technical issues. The authors thank A.V. Borisova for analytical discussions.

## Notes

https://github.com/il0106/Institute-of-Higher-Nervous-Activity-and-Neurophysiology

### Competing Interest Statement

The authors have declared no competing interest.

